# Enhanced Hydrogen Production Through Two-Stage Fermentation Coupling *Clostridium pasteurianum* and *Rhodobacter sphaeroides*

**DOI:** 10.64898/2025.12.14.694164

**Authors:** Huangwei Wang, Xin Zhang, Yufan Cheng, Xilin Ye, Jianhua Fan, Xu Zhang, LiXin Zhang, Gao-Yi Tan

## Abstract

Biological H_2_ production via dark fermentation is constrained by low yields and inhibitory metabolite accumulation. Coupling dark and photo fermentation effectively overcomes these issues by enhancing substrate utilization and reducing wastewater pollution, yet scalable systems for industrial application remain rare. This study presents a 10-L two-stage fermentation system (5L dark/5L photo bioreactors) using *Clostridium pasteurianum* DSM 525 and *Rhodobacter sphaeroides* ZX-5. Dark fermentation generated 3372 mL H_2_ from glucose, yielding effluent with 1.29 g/L acetic and 3.11 g/L butyric acids. After centrifugation, pH adjustment, and clinoptilolite deamination (≥60% efficiency), ammonium was reduced to <100 mg/L. The pretreated effluent, supplemented with concentrated RCVBN medium, served as photo-fermentation substrate. A butyric acid feeding strategy extended the process by 120 h, boosting H_2_ yield by 16.4%. The coupled system produced 6480 mL H_2_ (80% increase), degrading 50% acetic acid and 42% butyric acid. This work demonstrates a scalable bioreactor configuration integrating efficient biohydrogen production with value-added wastewater treatment for industrial bioenergy applications.

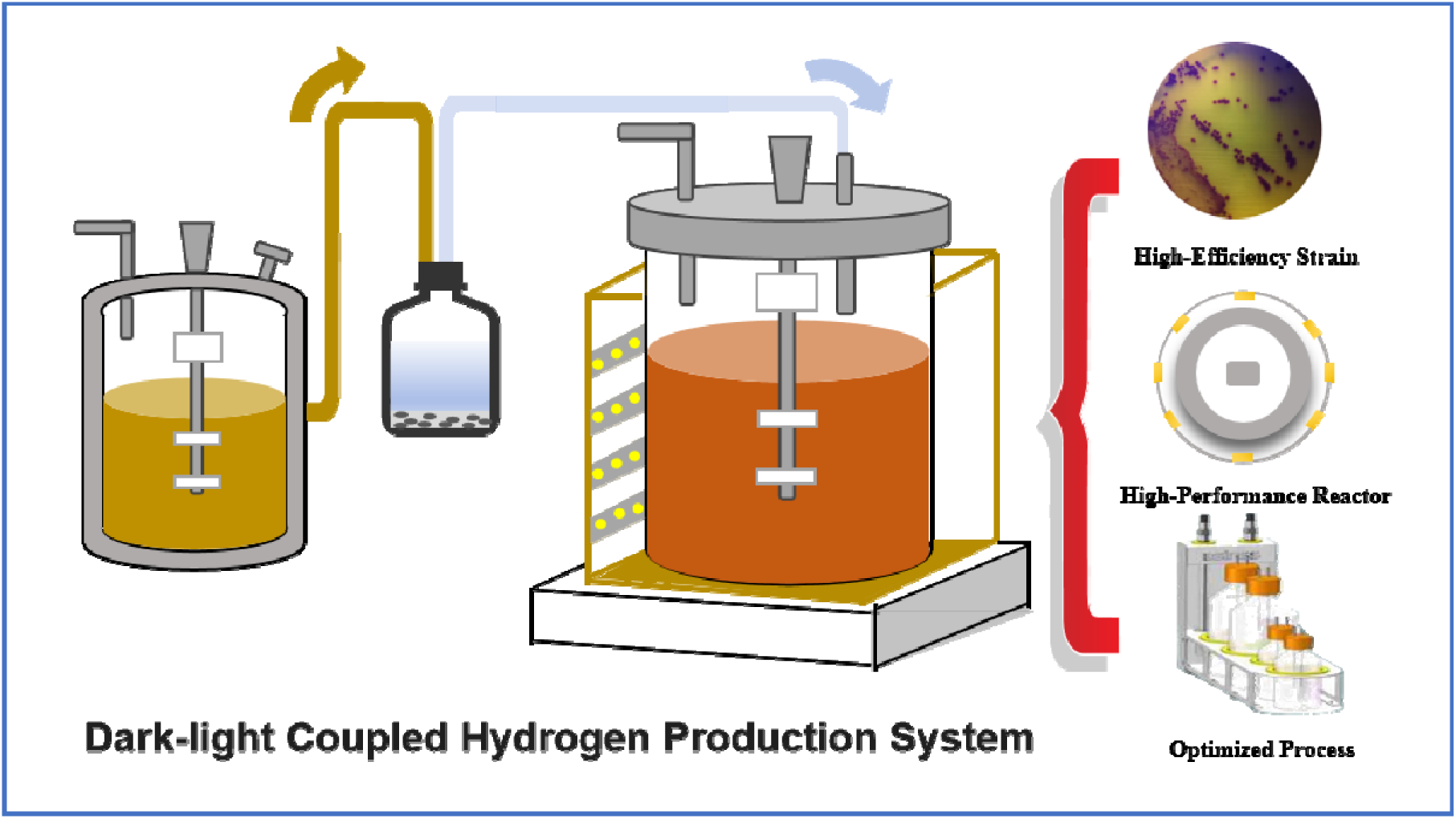
Graphic abstract.

## 1. Introduction

The global energy crisis, coupled with escalating environmental concerns, has highlighted the deficiencies of traditional H_2_ production methods. Most conventional techniques, such as steam methane reforming (Szablowski et al., 2025)and coal gasification (Duan et al., 2014), heavily rely on fossil fuels, resulting in substantial carbon emissions and a significant environmental impact. Although water electrolysis presents a cleaner alternative, its low energy efficiency and high operational costs restrict its widespread adoption (Xiuyuan et al., 2024). Consequently, the demand for sustainable and cost-effective alternatives has brought biological H_2_ production technologies into the spotlight. Biological H_2_ production, which utilizes microorganisms or enzymatic processes, harnesses renewable resources like organic waste and solar energy to generate H_2_. It offers environmental benefits and efficient resource utilization, opening a feasible new path for green H_2_ production.

Common biological hydrogen production processes include algal photosynthetic hydrogen production, anaerobic bacterial dark fermentation, and photosynthetic bacterial photo-fermentation. Single-stage biological H_2_ production processes encounter numerous challenges (Sagir et al., 2017). Firstly, while Dark Fermentation (DF) can efficiently produce H_2_, it results in a significant accumulation of by-products, such as acetic acid and butyric acid, during the process (Ergal et al., 2018). These acidic by-products can inhibit microbial activity in subsequent stages, thereby reducing overall H_2_ production efficiency. Secondly, Photo Fermentation (PF) relies on light energy to drive H_2_ production, but its efficiency is constrained by light intensity and bioreactor design, leading to a lower H_2_ yield compared to DF (Argun et al., 2008). Moreover, single-stage biological H_2_ processes often do not fully utilize the organic materials in the feedstock, resulting in inefficient resource utilization and limiting economic viability and industrial application. In the field of biological H_2_ production, two-stage coupled processes have been attracting attention. This involves combining DF with PF or biological photolysis to enhance H_2_ production while reducing the Chemical Oxygen Demand (COD) of wastewater, achieving green emissions. Two-stage coupled H_2_ production can decrease environmental pollution and enable resource recycling, offering an effective solution for sustainable green H_2_ production.

In recent years, studies have shown that two-stage coupling not only increases biological hydrogen production but also enhances the energy conversion efficiency of bioenergy systems (Huang et al., 2020; Morsy et al., 2019; Rezaeitavabe et al., 2020). For example, Das et al. optimized the system by controlling pH, allowing the PF stage to efficiently convert organic acids from DF effluent, achieving 650 mL H□/L (212.4 mL H□/g TVFA) (Das & Basak, 2025). DF/PF coupling technology faces multiple challenges during scale-up. The DF/PF system typically requires multi-stage bioreactors. For the dark fermentation stage, conventional fermenters are commonly used for hydrogen production (Teshnizi et al., 2025; Tun et al., 2025). In contrast, the photo-fermentation stage often employs tubular photobioreactors (Grazia et al., 2023; Tian et al., 2021) in batch mode to enhance light efficiency. However, due to reactor limitations, a single photobioreactor alone is unlikely to further improve hydrogen yield through process optimization. Moreover, coupling dark and photo-fermentation at different reactor scales presents significant challenges. Therefore, achieving efficient dark-light coupling at the reactor scale through advanced bioreactor design and photo-fermentation process optimization is key to overcoming the current limitations of hydrogen productivity in DF/PF coupled systems.

In this study, the dark fermentation strain utilized is *Clostridium pasteurianum* (*C. pasteurianum*) DSM 525, one of the highest H_2_-producing dark fermentation bacteria currently known (Poehlein et al., 2015). The photosynthetic fermentation strain employed is *Rhodobacter sphaeroides* (*R. sphaeroides*) ZX-5, which can utilize various organic acids, such as butyrate and acetate, for H_2_ production (Tao et al., 2008) and exhibits higher H_2_ yields compared to other photosynthetic bacteria. In the coupled process, we have adopted an innovative photobioreactor design and a butyrate-coupling feeding strategy to enhance the integration of the dark fermentation system with the photo-fermentation system, improving H_2_ yield at a 5L bioreactor scale. This work not only advances our understanding of dark-light coupled fermentation but also provides valuable insights for optimizing biological H_2_ production in industrial applications.

## 2. Materials and methods

### 2.1. Cultivation conditions of *R*. *sphaeroides* ZX-5

*R. sphaeroides ZX-5* was obtained from the Institute of Ecology and Soil, Guangdong Academy of Sciences, cultured in LB medium(add 1% proteose, 1% sodium chloride and 0.5% yeast extract to H_2_O and make up to 1000 mL, stir evenly). This was incubated in a shaker at 37°C and 220 rpm for 24 hours. The pre-cultured suspension (viable count ≥1×10□ CFU/mL) was inoculated aseptically at 3.3% (v/v) into a sterilized 50 mL conical flask fermentation system containing 40 mL of RCVBN medium (Table S1) previously sterilized at 115°C for 30 min. The system was sealed using a reverse-punctured silicone stopper. Gas storage was achieved using a 20 mL medical syringe. This bioreactor system was placed in a photo-shaker incubator set at 180 rpm, a temperature gradient of 33.0 ± 2.0°C, and continuous light irradiation within the wavelength range of 400-700 nm at a photon flux density of 120 ± 5 μmol·m□²·s□¹. The cultivation period was dynamically adjusted based on gas chromatography (GC) monitoring values.

### 2.2. Cultivation conditions of *C. pasteurianum* DSM 525

*C. pasteurianum* DSM 525 was donated by Yantai Coastal Zone Research Institute, Chinese Academy of Sciences. Glycerol stock stored at 4°C was taken. A small amount of bacterial suspension (sampled by inserting a syringe needle into the stopper, inverting the bottle, drawing liquid into the syringe, and removing the syringe) was transferred into basal medium. Cultivation was carried out in 100 mL penicillin bottles containing 40 mL of medium, incubated statically at 37°C for 12 h. For scale-up: After the culture became turbid, 5% of the culture was aseptically transferred using a 5 mL syringe into MSG liquid medium (Table S2) in a 250 mL penicillin bottle containing 100 mL of medium. This was placed in a shaker incubator at 35°C and 100 rpm. When the culture color changed from pale yellow to milky white and the rubber stopper was pushed up, samples could be taken to measure OD_600_.

### 2.3. Bioreactor fermentation of *R. sphaeroides* ZX-5

Fifty well-rounded, uniformly sized, and evenly distributed *R. sphaeroides* ZX-5 single colonies grown for 6-7 days were picked and added to 200 mL of LB medium. This was incubated in a shaker at 37°C and 220 rpm for 24 hours to serve as the seed culture. A 5L bioreactor equipped with an orbitally arranged light strip providing an illumination intensity of 1.8 W/m²/nm (at a light path of 3 cm) was used. The bioreactor contained 3L of RCVBN medium sterilized at 115°C for 30 min. Inoculation was performed at 5% (v/v). The pH during fermentation was controlled at 7.5 ± 0.3 using a phosphate buffer. A facultative culture process was employed without additional oxygen supplementation. The produced gas was collected using the water displacement method, and the H_2_ component in the mixed gas was determined by GC.

### 2.4. Bioreactor fermentation of *C. pasteurianum* DSM 525

The fermenter was cleaned and sterilized empty beforehand. After empty sterilization, a 5x concentrated liquid medium solution was prepared according to the formula and added to the vessel along with approximately 1.5 L of H□O. The pH of the medium was adjusted to 6.6 (initial pH ∼7.1) before sterilization. After sterilization, the vessel temperature was set to 35°C, and nitrogen gas was continuously sparged into the vessel for about 1.5 hours. After the 1.5 h nitrogen purge, the gas flow was stopped, and inoculation was performed under a flame. During inoculation, the seed culture (5% inoculum, ∼0.1 L), glucose solution (∼0.1 L), and a mixture of trace elements, mineral solution, and vitamin solution (prepared as 5x concentrate, ∼0.1 L) were sequentially added into the fermenter within a short period to achieve a total fermentation volume of approximately 2 L. After inoculation, nitrogen gas was sparged again for about 30 min, then both inlet and outlet gas ports were closed. The produced gas was collected using the water displacement method, and the H_2_ component in the mixed gas was determined by GC.

### 2.5. Analytical methods

pH was measured with a Mettler laboratory pH meter, light intensity was measured with a Delixi digital light meter, and biomass concentration (cell density) was determined using a UV spectrophotometer. For biomass determination, 0.5 mL of culture broth was transferred to a 2 mL centrifuge tube, diluted to 2 mL with H□O, and the OD□□□ was recorded.

### 2.6. Mixed gas composition and content determination

Mixed gas produced during fermentation was collected using a 50 mL syringe. Gas composition and content were analyzed using a Gas Chromatograph (Model GC8950, provided by Shanghai Tianpu Analytical Instrument Co., Ltd.). The instrument used a TCD detector and a TDX-01 chromatographic column, with high-purity argon as the carrier gas. During detection, the injector, column oven, and detector temperatures were all set to 100°C. After system temperature stabilized, the bridge current was adjusted to 50 mA. The injection volume was set to 1 mL of mixed gas. A standard curve was established using high-purity H_2_, and the H_2_ content was calculated using the external standard method (specific curve shown in Fig. S1).

### 2.7. Residual acid composition and content determination in fermentation broth

Residual acids in the fermentation broth were determined by HPLC. Column: Hypersil ODS-100s C18, 5 μm, 4.6×250 mm. Mobile phase: 5 mM H□SO□, flow rate: 0.8 mL/min. Detection wavelength: 210 nm, temperature: 25°C. A volume of fermentation broth was centrifuged at 12,000 rpm for 10 min to remove cells and other insoluble matter. Then, 1 mL of the supernatant was filtered through a 0.22 μm membrane into an HPLC vial for analysis. The standard curve was determined by preparing standard solutions of acetic acid/butyric acid/glucose at concentrations of 1, 3, 10, 15, and 30 mM. Under the same chromatographic conditions (injection volume 20 μL), a plot of chromatographic peak area versus standard concentration was generated. Fermentation broth samples were analyzed under the same LC conditions. The peak areas were calculated using the chromatography workstation software, and the concentrations of acetic acid, butyric acid, and glucose in the broth were determined from the standard curves (Fig. S2).

### 2.8. Residual ammonium nitrogen content determination in fermentation broth

Ammonium nitrogen concentration was determined using Nessler’s reagent spectrophotometry. First, a standard curve was prepared(Fig. S3): Take 0.00∼10.00 mL of ammonium standard solution (0.010 mg/mL NH□^+^-N) into 50 mL reaction tubes, bring to volume with water, then sequentially add 1.0 mL of potassium sodium tartrate (500 g/L) and 1.0 mL of Nessler’s reagent. Vortex mix (640 rad/min × 60 s), develop color in the dark at 25 ± 1°C for 15 min, and measure absorbance at 420 nm to establish a linear regression model. For sample measurement: Take 2.00 mL of fermentation broth, dilute to 50 mL, add the color reagents following the same method, vortex mix (2000 rpm × 30 s), develop color in the dark at 25 ± 1°C for 10 min, then measure absorbance. The ammonium nitrogen concentration was finally calculated using the formula:

Ammonium nitrogen concentration (mg/L) = (Ammonium nitrogen mass obtained from the standard curve (μg) / 2.00 mL) × 1000

(Note: Samples were diluted 25-fold; results need correction for the dilution factor).

### 2.9. Calculation of parameters related to H_2_ production process

(1) H_2_ yield formula:

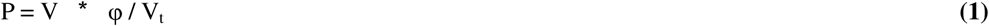

Where P represents hydrogen yield (mL/L), V represents the volume of mixed gas produced (mL), φ represents hydrogen content (%), and V_t_ represents the volume of medium in the hydrogen production system (mL).
(2) Average H_2_ production rate formula:

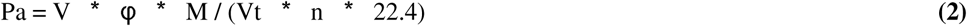

Where P_a_ represents the average H_2_ production rate (g/L/day), V represents the volume of mixed gas produced (L), φ represents H_2_ content (%), V_t_ represents the volume of medium in the H_2_ production system (L), n represents the cultivation period (days), M is the molar mass of H_2_ (2 g/mol), and 22.4 L/mol is the molar volume at STP.
(3) Substrate conversion efficiency formula:

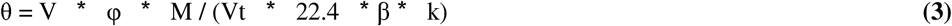

Where θ represents substrate conversion efficiency (%), V represents the volume of mixed gas produced (L), φ represents H_2_ content (%), k represents the substrate concentration (g/L), V_t_ represents the volume of medium in the H_2_ production system (L), M represents the relative molecular mass of the substrate (g/mol), and β represents the theoretical stoichiometric ratio of H□ produced per mole of substrate completely converted. (Note: The formula structure aims to calculate the percentage of the theoretical maximum H_2_ yield achieved from the consumed substrate. The term (22.4 / M) * 1000 converts substrate mass to theoretical moles of H□ based on β).

## 3. Results and discussion

### 3.1. Study on the Adaptability of *R. sphaeroides* ZX-5 to Wastewater

**Fig 1**a showed H_2_ production results from dark fermentation of *C. pasteurianum* DSM 525 in 100 mL serum bottles, which exhibited rapid H_2_ accumulation, reaching 180 mL within 40 hours (80% of total yield). Production plateaued abruptly after 60 hours, indicating substrate depletion or metabolic shift. In contrast, **Fig 1**b showed H_2_ production results from photo-fermentation of *R. sphaeroides* ZX-5 in 50 mL conical flasks, sustained production over 96 hours, achieving 75 mL without plateauing.

**Fig 1.**
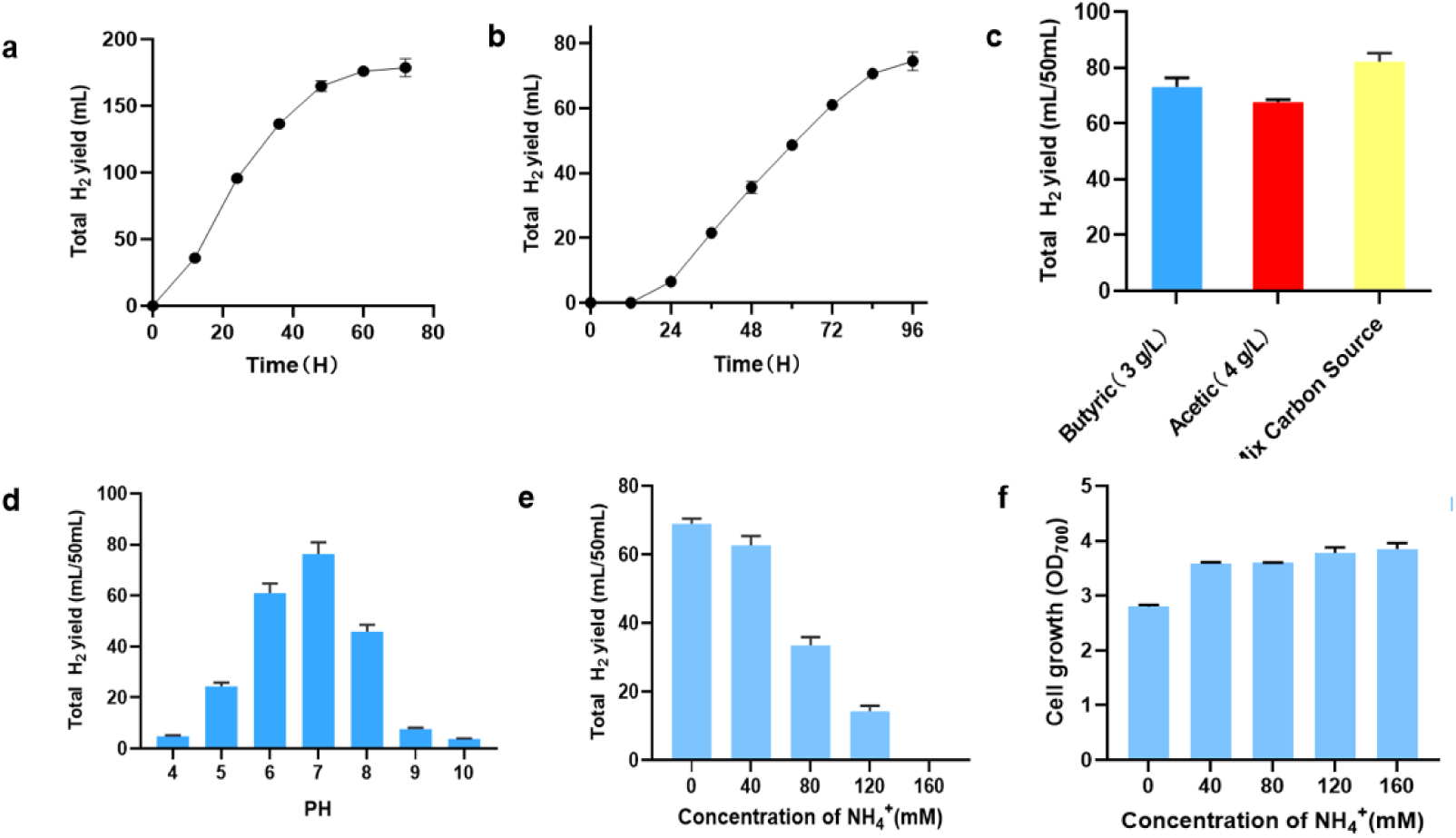
Effects of major components in wastewater on H_2_ production. (a) H_2_ production results from dark fermentation of *C. pasteurianum* DSM 525 in 100 mL serum bottles. (b) H_2_ production results from photo-fermentation of *R. sphaeroides* ZX-5 in 50 mL conical flasks. (c) Influence of major carbon sources in wastewater on *R. sphaeroides* ZX-5 hydrogen production. The mixed carbon sources consist of butyric acid (2.964 g/L), acetic acid (1.28 g/L), and glucose (1.2 g/L). (d) Effect of pH on *R. sphaeroides* ZX-5 hydrogen production (ranging from 4.0 to 10.0). (e) Impact of NH_4_ ^+^ concentration on H_2_ production of *R. sphaeroides* ZX-5. (f) Impact of NH_4_ ^+^ concentration on cell growth of *R. sphaeroides* ZX-5. All strains were cultivated in a shaking incubator at 32 °C and 180 rpm for 120 h under 7000 lux light, with a 50 mL fermentation system.

To investigate the adaptability of *R. sphaeroides* ZX-5 to Clostridium wastewater, the components of the wastewater were determined after 48 hours of dark fermentation, as shown in Table S3. The wastewater was dominated by volatile fatty acids (VFAs), primarily butyric acid (2.964 g/L) and acetic acid (1.28 g/L), which accounted for 99.3% of the total VFAs. Additionally, the wastewater contained high nitrogen sources, including 6 mg/L ammonium nitrogen and 144 mg/L total nitrogen, likely derived from decomposed bacterial proteins and potentially affecting microbial physicochemical activity. A notable feature was the high chemical oxygen demand (COD, 10190 mg/L), which might lead to excessively low oxidation-reduction potential (ORP), significantly impacting process efficiency. The wastewater also contained 2700 mg/L Na^+^ (0.12 M). Studies have shown that Na□ neutralizes the negative charges on the lipid headgroups of cell membranes, forming an “ion-screening” effect that significantly enhances membrane structural stability and maintains transmembrane proton gradients. For instance, Schoepp reported that 0.1 M Na^+^ increased the photosynthetic electron transport rate of *Chromatium vinosum* to 2.3 times that under sodium-free conditions(Michael et al., 2013). Therefore, desalination of the wastewater was not considered in this study. Overall, high nitrogen content and acidity are the primary characteristics of the wastewater, representing key challenges for H_2_ production in subsequent coupled processes.

To determine the major components in wastewater affecting hydrogen (H_2)_ production by *R. sphaeroides* ZX-5, the individual effects of mixed carbon sources, pH, and ammonium concentration were investigated. Considering that the optimal carbon source concentrations for *R. sphaeroides* ZX-5 are acetic acid (4 g/L) and butyric acid (3 g/L) (Li et al., 2009), the substrate conditions were simulated to mimic the mixed carbon sources found in wastewater, including acetic acid (1.2 g/L), butyric acid (2.94 g/L), and glucose (1.2 g/L) for H_2_ production fermentation(**Fig 1c**). Experimental results indicated that during photo-fermentative H_2_ production using mixed carbon sources, *R. sphaeroides* ZX-5 preferentially utilized glucose as an energy source to accelerate cell proliferation, resulting in low H_2_ production in the early stage. In the middle stage, H_2_ production efficiency increased significantly, mainly driven by energy carriers (such as reduced coenzymes) generated from the catabolism of butyric and acetic acids, which promoted substantial H_2_ generation. In the late stage, the H_2_ production rate slowed down due to the accumulation of residual butyric acid and the depletion of other substrates. Compared with single-substrate systems, mixed carbon sources enhanced H_2_ production (an 11.36% increase) and cell growth (a 14.66% increase), leveraging the metabolic preferences of individual carbon sources while avoiding their drawbacks, thus generally facilitating photo-fermentative H_2_ production by *R. sphaeroides* ZX-5. This suggests that mixed carbon sources are not the primary limiting factor inhibiting H_2_ production from wastewater.

The pH value is crucial for the growth, metabolism, and survival of bacteria. The optimal pH for photosynthetic bacteria is generally around 7.0 (Cheng et al., 2011). As shown in **Fig 1d**, H_2_ production peaked at pH 7.0, approximately 15-fold higher than under acidic conditions. H_2_ yield increased significantly with pH elevation in the range of 4-7, but decreased sharply at pH > 7. It is hypothesized that under acidic conditions (pH < 6), proton backflow disrupts transmembrane gradients and inhibits ATP synthesis, while alkaline environments (pH > 8) reduce the supply efficiency of reduced coenzymes (NAD(P)H) by altering membrane lipid structures. Collectively, pH 7.0 represents the optimal condition for H_2_ production by *R. sphaeroides* ZX-5. In subsequent treatment processes, adjusting wastewater to neutrality and adding phosphate buffer (0.2 M, pH=7) can mitigate the adverse effects of pH on H_2_ production.

Nitrogen is a critical nutritional element for bacterial growth, directly influencing their metabolism and reproduction. In addition to utilizing nitrogen sources for growth, *R. sphaeroides* ZX-5 can also reduce N_2_ to NH_3_ using nitrogenase (Seefeldt et al., 2012). If the catalytic mechanism of nitrogenase is considered a normal chemical reaction, theoretically, the continuously generated ammonium salts would impede the reaction from proceeding forward, leading to a gradual decline in H_2_ production. As depicted in **Fig 1d**, *R. sphaeroides* ZX-5 exhibited the highest H_2_ production (72.36 mL) under ammonium-free conditions. When the ammonium concentration was 40 mg/L, the optical density at 700 nm (OD_700_) increased significantly by 23% (**Fig 1e**), while H_2_ production decreased by 21% (**Fig 1f**), indicating that sufficient nitrogen sources prioritize ammonium metabolism, which consumes intracellular ATP and reducing power, thereby inhibiting the hydrogen-producing metabolic flux. As ammonium concentration increased to 160 mg/L, bacterial biomass continued to rise while hydrogen production dropped sharply. Hydrogen production system collapsed entirely at ammonium concentrations ≥120 mg/L. This occurs because excessive NH_4_ ^+^ directly inhibits nitrogenase activity via substrate effects and may disrupt hydrogen production pathways by regulating nitrogenase-related gene expression. With ammonium concentration in wastewater reaching 144 mg/L(significantly higher than the optimal 40 mg/L), ammonium toxicity is identified as the primary limiting factor for H_2_ production by *R. sphaeroides* ZX-5 using waste acids.

### 3.2. Pilot-scale test of Clostridium wastewater treatment process

It is established that the primary solid matter in Clostridium wastewater consists of spent microbial biomass, typically removed via centrifugation. Some studies additionally employ autoclaving to eliminate contaminants and colloidal substances that may impede light penetration. Following centrifugation at 12,000×g RCF for 10 min, the wastewater pH was adjusted to 7.0 ± 0.21 using 0.2 M NaOH, followed by supplementation with 5% (v/v) of 0.2 M phosphate buffer (pH 7.0). To remove nitrogen sources, clinoptilolite (50-mesh) was utilized for pretreating dark fermentation effluent (DFE) as illustrated in **Fig 2b**. Zeolites are natural or synthetic microporous aluminosilicate minerals characterized by well-defined three-dimensional crystalline structures containing abundant microchannels and cavities. Their general chemical formula is M_x_/_n_[(AlO_2_)_x_(SiO_2_)_y_]_z_H_2_O, where M denotes exchangeable cations (e.g., Na^+^□K^+^□Ca^2+^). Zeolites remove ammonium ions (NH_4_^+^) through ion exchange and physical adsorption. When ammonium-containing water permeates zeolites, NH_4_^+^(diameter ≈ 2.86 Å) effectively enters pores (typically 3-10 Å) and undergoes cation exchange, thereby immobilizing NH_4_^+^.

**Fig 2.**
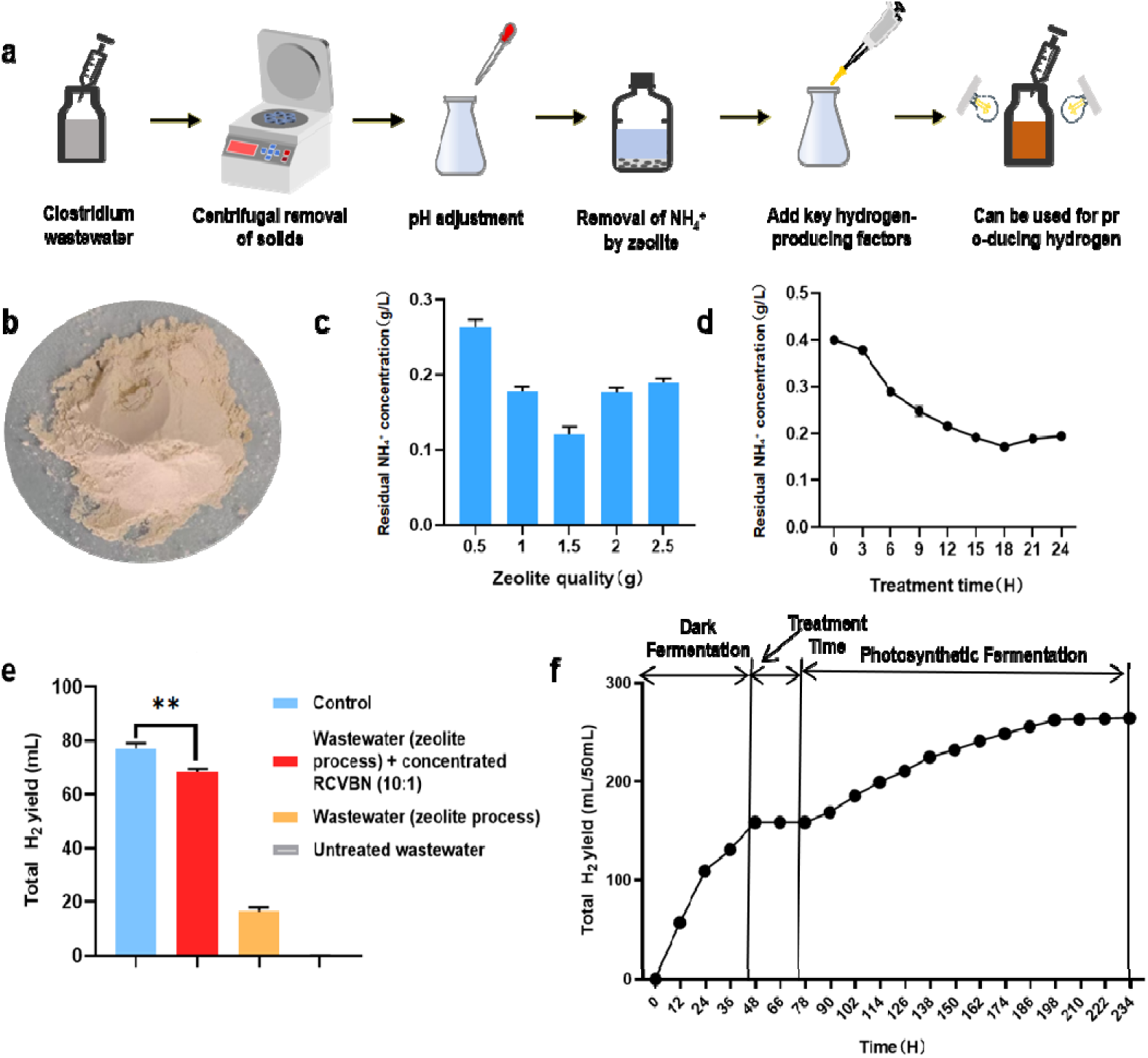
Process for utilizing Clostridium wastewater in photosynthetic H_2_ production based on zeolite. (a) Pretreatment process flow of Clostridium wastewater, including centrifugation for solid removal, pH adjustment, zeolite treatment, and supplementation of key H_2_-producing factors. (b) Clinoptilolite used for ammonium salt removal in wastewater. (c) Effect of zeolite dosage on ammonium removal efficiency. (d) Effect of treatment duration on ammonium removal efficiency. (e) Impact of different wastewater treatment processes on H_2_ production of *R. sphaeroides* ZX-5. (f) Changes in total H_2_ accumulation during coupled photo-dark fermentation of wastewater in a 50 mL system.

To determine the removal efficiency of NH_4_^+^ from Clostridium wastewater, the effects of zeolite dosage and treatment duration were investigated. As illustrated in **Fig 2c**, ammonium adsorption exhibited significant nonlinearity with increasing clinoptilolite dosage. The removal efficiency increased from 32.5% to 65% when the dosage rose from 0.5 g to 1.5 g, but it declined by approximately 20% beyond 1.5 g due to pore blockage from particle aggregation, reducing the effective surface area utilization. Thus, the optimal adsorption occurred at 1.5 g. **Fig 2d** reveals that NH_4_^+^ adsorption progressed through three phases: rapid adsorption (0-15 hours), equilibrium (15-18 hours), and desorption (>18 hours). During the first 18 hours, the NH_4_^+^ concentration decreased from 0.4 g/L to 0.172 g/L (57% removal), following pseudo-second-order kinetics, indicating that ion exchange and chemisorption dominated NH_4_^+^ fixation. The anomalous increase observed after 18 hours is attributed to the partial release of NH_4_^+^ due to electrostatic repulsion upon saturation. The maximum adsorption capacity (0.228 g/g) was achieved at 18 hours, indicating that optimal conditions are 1.5 g of clinoptilolite per 10 mL of wastewater with an 18-hour treatment period.

The post-treatment NH_4_^+^ concentration was 0.098 ± 0.0058 g/L. Organic acid levels also decreased: acetic acid to 0.98 ± 0.12 g/L and butyric acid to 2.78 ± 0.07 g/L. When zeolite-treated effluent was utilized (**Fig 2e**), the H_2_ yield was only 23.4% of the control, which may be attributed to the lack of some essential key components, contains mineral salts, vitamins, metal ions (e.g., Fe^2+^□Mo^6+^ for hydrogenase activation), and cofactors whose absence limits hydrogenase synthesis and electron transfer (Seefeldt et al., 2012). These key components are all included in the RCVBN photosynthetic hydrogen production medium. Therefore, it can be used as a nutritional supplement solution by mixing it with wastewater, and it is speculated that this could restore the hydrogen production activity of the wastewater. Testing showed that 10× RCVBN (without carborn) maximized retention of native carbon sources while minimizing costs. Subsequent cofactor supplementation enabled efficient organic acid utilization by *R. sphaeroides* ZX-5 (**Fig 2e**).

The integrated process (**Fig 2a**) involves: (1) centrifugation and pH adjustment; (2) clinoptilolite treatment (1.5 g/10 mL, 18 h); (3) mixing treated wastewater with 10× RCVBN at 10:1 ratio for *R. sphaeroides* ZX-5 cultivation. In 50 mL coupled systems (**Fig 2f**), dark fermentation yielded 183.56 ± 1.36 mL H_2_ from wastewater containing butyric acid (2.88 ± 0.14 g/L) and acetic acid (1.212 ± 0.21 g/L). After treatment, 5% (v/v) inoculum of *R. sphaeroides* ZX-5 was cultured at 8000 lux, 32°C, and 180 rpm for 120 h. Total H_2_ production reached 274.66 ± 2.27 mL, with 66.77% acetic acid and 70.66% butyric acid removal, demonstrating successful process integration.

### 3.3. Experiment on photo-dark coupled H_2_ production process in 5-L reactor

#### 3.3.1. Scale amplification investigation of dark fermentation bioreactors

To achieve bioreactor-scale amplification of the photo-dark coupled fermentation process, the dark fermentative H_2_ production by *C. pasteurianum* DSM 525 in a 5L bioreactor was first investigated(**Fig 3b**). As depicted in **Fig 3a**, the 3-L cultivation system (initial glucose: 10 g/L, initial pH: 6.6) exhibited distinct phase-specific dynamics in H_2_ production and metabolic profiles. During 0-48 h, H_2_ accumulation demonstrated an exponential increase phase, reaching a peak of 3372.4 mL, indicating completion of the primary H_2_ genic stage within 48 h. Correspondingly, biomass (OD_600_) peaked between 24-36 h, entering a decay phase post-48 h. Butyric and acetic acids accumulated progressively throughout fermentation, attaining final concentrations of 2.95 g/L (**Fig 3c**) and 1.3 g/L (**Fig 3d**), respectively. Notably, butyrate accumulation accelerated more rapidly, achieving 2.95 g/L by 48 h.

**Fig 3.**
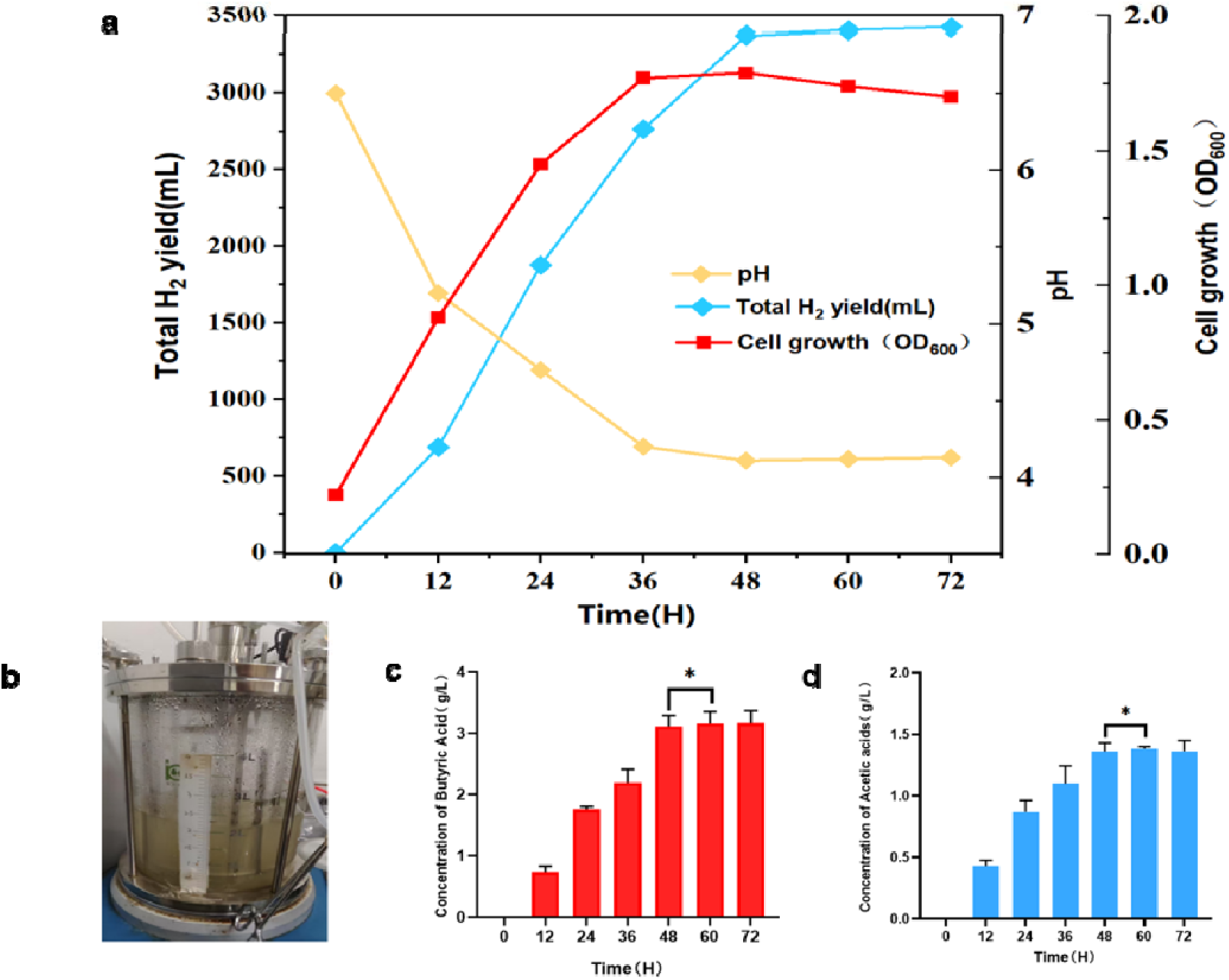
H_2_ Production by *C. pasteurianum* DSM 525 in a 5L bioreactor via dark fermentation. (a) Temporal profiles of pH, biomass growth (OD_600_), and cumulative H_2_ production. (b) H_2_ production by dark fermentation in a 5L bioreactor. (c) Butyric acid accumulation dynamics (g/L) at indicated time points. (d) Acetic acid accumulation dynamics (g/L) at indicated time points. Strains were cultured anaerobically for 72 h under dark fermentation conditions.

The 48-h timepoint was selected as the optimal coupling node for the dark-photo fermentation system based on three critical factors: (a) H_2_ yield saturation: 97.4% of total H_2_ production was completed by 48 h, with extended dark fermentation contributing <5% additional yield. (b) Ideal substrate thresholds: Butyrate (2.95 g/L) and acetate (1.3 g/L) concentrations approached the optimal range (<3.0 g/L) for subsequent photo-fermentation without significant further accumulation. (c) Metabolic inhibition risk: Initiation of cell lysis beyond 48 h may release inhibitory byproducts detrimental to photosynthetic H_2_ production.

#### 3.3.2. Investigation of photo fermentative H_2_ production process in a 5L bioreactor

This study examined the photo-fermentative H_2_ production process by *R. sphaeroides* ZX-5 in a 5-L bioreactor. Unlike traditional bioreactor processes, the illumination conditions are the most critical factor in photosynthetic fermentation. The photobioreactor used in this study demonstrated light-shielding effects—a phenomenon where uneven light distribution, caused by high cell density or bacterial aggregation, creates dark zones that significantly reduce H_2_ production efficiency. This effect mainly stems from excessive light absorption by surface-layer bacteria, cell aggregation, and light scattering due to broth turbidity, which collectively prevent deep-layer cells from performing efficient photophosphorylation and suppress nitrogenase activity. Such dark zones not only compel bacteria to follow non-hydrogenic metabolic pathways but also decrease overall H_2_ evolution rates.

To mitigate the adverse impacts of light-shielding effects, three illumination configurations were tested: Inner Wall Illumination, Axial Illumination, and Outer Track Illumination (**Fig 4a**). The first two are commercially available technologies using incandescent tubes embedded in the bioreactor with fixed maximum light intensity (15000 lux), providing localized illumination. These systems face limitations in both maximum irradiance and insufficient radiation coverage for outer-layer cells. Building upon these, we designed a novel Outer Track Illumination system (Fig. S4) employing point-source LED beads (max 40000 lux). This customizable external orbital system allows flexible selection and replacement of light strips while enabling denser illumination arrangements, reducing dark zones by 20-30% compared to conventional systems (Biology et al., 2019).

**Fig 4.**
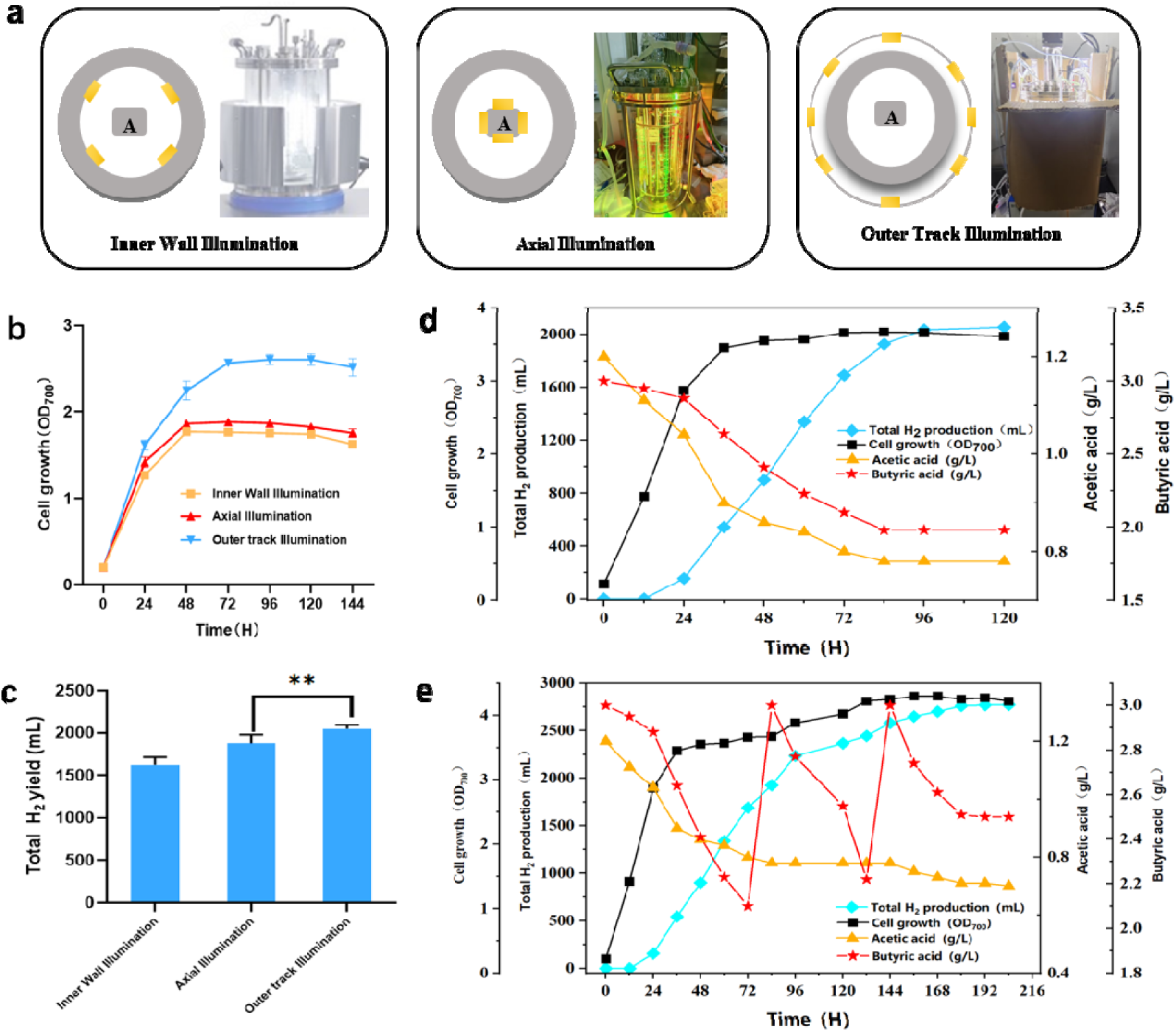
Results of photo-fermentative biohydrogen production in a 5L bioreactor. (a) Schematics of three photobioreactor types: Inner Wall Illumination, Axial Illumination, and Outer Track Illumination. (b) Biomass growth profiles of *Rhodobacter sphaeroides* ZX-5 across photobioreactor configurations. (c) Comparative H_2_ production yields under respective illumination systems. All strains were cultivated in a 5L bioreactor at 32 °C and 180 rpm for 120 h. (d) Temporal profiles of key parameters during a 120-h batch fermentation of photo-fermentative biohydrogen production in a 5L photobioreactor. (e) Temporal profiles of key parameters during a 220-h fed-batch fermentation of photo-fermentative biohydrogen production in a 5L photobioreactor. At 84 h, a total of 100 mL of 90 g/L butyrate solution was continuously supplemented, restoring the butyrate concentration to 3.0 g/L. Detection of butyrate concentration below 2 g/L at 144 h prompted a second supplementation of 91.66 mL of 90 g/L butyrate solution, restoring the butyrate concentration to 3.0 g/L.

Outer Track Illumination significantly outperformed Inner Wall and Axial Illumination in both cell growth and H_2_ production. Biomass accumulation curves (**Fig 4b**) showed that Outer Track maintained the highest OD_700_ (peak 2.8) throughout 0-144 h, exceeding Inner Wall and Axial systems by 27% and 40%, respectively. H_2_ production(**Fig 4c**) reached 2340 mL under Outer Track—44% and 48% higher than Inner Wall (1620 mL) and Axial (1580 mL) configurations. The uniform spatial distribution achieved by the 8-point annular layout effectively expanded the illuminated surface area and minimized dark zones, promoting efficient metabolic activity throughout the fermentation cycle. In contrast, Inner Wall Illumination (4-point inner layout) suffered from limited coverage, while Axial Illumination’s central radiation pattern resulted in insufficient penetration depth to activate peripheral cells.

The photo-fermentative H_2_ production process by *R. sphaeroides* ZX-5 in a 5L bioreactor was simulated using substrate concentrations representative of Clostridium wastewater effluent (butyrate: 2.88 g/L, acetate: 1.212 g/L) as the initial substrate levels (**Fig 4d**, Fig. S5). H_2_ production exhibited a significant increase during the initial 80 hours of fermentation, indicating sufficient carbon source availability and robust microbial metabolic activity in this phase. However, the H_2_ production rate declined sharply beyond 80 hours (approximately a 53% reduction), presumably due to metabolic limitations necessitating substrate replenishment via feeding. The inflection points in H_2_ production (84 hours) correlated strongly with the onset of accelerated butyrate consumption, identifying butyrate as the key carbon source limiting H_2_ genesis. Based on these kinetic profiles, a feeding strategy was devised to initiate substrate supplementation at 84 hours, coinciding with near-depletion, to sustain metabolic flux and prolong the photo-fermentative H_2_ production process.

Following the determination of the optimal feeding timepoint (84 h), the effect of supplement concentration on H_2_ production was investigated. At 84 h, 100 mL of concentrated medium containing butyrate at concentrations of 45, 90, 135, or 180 g/L was introduced into the fermentation system. This corresponded to calculated final butyrate concentration increases in the 3.5 L working volume of 1.29, 2.57, 3.86, 5.14, and 6.43 g/L, respectively. Total H_2_ production was monitored throughout the fermentation (Fig. S6). Systematic investigation of the butyrate feeding concentration revealed that 90 g/L (yielding a final concentration increase of 2.57 g/L) constituted the distinct optimum. This concentration maximized H_2_ production (2500 mL), representing a 25% increase over the unfed control (2000 mL). Lower concentration supplementation (45 g/L, +1.29 g/L) yielded only marginal improvement (2200 mL, +17%). Conversely, concentrations exceeding 90 g/L (135-180 g/L, +3.86-5.14 g/L) induced progressive inhibition, manifested as reduced H_2_ yields attributable to acid stress and metabolic imbalance. These results confirm that precise substrate feeding at 84 h balances carbon flux, maximizes nitrogenase activity while avoiding catabolite repression, thereby establishing 90 g/L as the critical threshold for the optimized fed-batch photo-fermentation process.

For process validation in the 5L bioreactor, a 90 g/L butyrate solution (300 mL reservoir) was used as the feedstock. At 84 hours, a total of 100 mL of this solution was continuously fed, restoring the butyrate concentration to 3.0 g/L. Subsequently, residual substrate concentrations were continuously monitored, and butyrate levels were maintained at approximately ≥ 2.0 g/L until the fermentation endpoint, defined as the point where H_2_ production ceased to change significantly over a 48-h period. The temporal profiles of key parameters—including cumulative H_2_ production, cell density (OD_700_), and residual acetate and butyrate concentrations in the medium—during this fed-batch operation are presented. **Fig 4e**.

The implemented butyrate feeding strategy in the 5L bioreactor effectively alleviated substrate limitation. This droved a sustained increase in H_2_ production from 1925.28 mL at 84 h to 2770.6 mL at 204 h, representing a total increase of 44% and significantly extending the H_2_ production period. Concurrently, robust cell viability was maintained (OD_700_ increased from 3.67 to 4.22), butyrate concentration fluctuated within a controlled range (2.1-3.0 g/L), and acetate was continuously consumed, collectively indicating efficient channeling of carbon metabolism towards H_2_ production. H_2_ production stabilized by 204 h (<0.1% change over 48 h), marking the fermentation endpoint. This feeding protocol, by precisely maintaining substrate concentration, achieved a remarkable extension of the H_2_ production phase by over 120 h and a 44% enhancement in total yield. However, residual acetate and butyrate were still present at the endpoint. This incomplete substrate utilization may be attributed to the gradual entry of the microbial population into the decay phase following the secondary feeding. Further process optimization studies could explore reducing the number of feeding points to promote more complete consumption of residual substrates within the system.

#### 3.3.3. Pilot-scale demonstration of the photo-dark coupled H_2_ production process in 5L bioreactors

To establish a light-dark two-stage coupled H_2_ production process, research on dark fermentation and photo-fermentation processes was conducted in a 5L bioreactor. The two fermentation stages were coupled through feeding, as depicted in Figure **Fig 5a**, where the 5L reaction flask contained 100 g of sterilized zeolite. Initially, dark fermentation for H_2_ production was performed in a 5L bioreactor (with a 2L reaction system) over a 48-hour fermentation cycle. At the cycle’s conclusion, the pH was measured at 4.2, the acetate concentration was 1.29 g/L, and the butyrate concentration was 3.11 g/L. The resulting 2L dark fermentation wastewater was then transferred to a 5L reaction flask, and the pH was adjusted to 6.5-6.7 using a 0.2 M NaOH solution added dropwise with a rubber-tipped dropper. The wastewater remained in the reaction flask for 18 hours until the ammonium salt concentration decreased by 60% or more, after which it was pumped into a 5L photobioreactor. The photobioreactor already held 0.3 L of pre-sterilized 10-fold concentrated RCVBN medium, which was thoroughly mixed with the treated 3 L of Clostridium waste liquid, resulting in a total volume of 3.3 L and diluting the RCVBN medium to a 1-fold concentration. The photosynthetic bacteria medium contained 1.2 g/L of acetic acid and 3 g/L of butyric acid and was inoculated at a rate of 5%. The variations in each parameter throughout the fermentation process are illustrated in **Fig 5b**.

**Fig 5.**
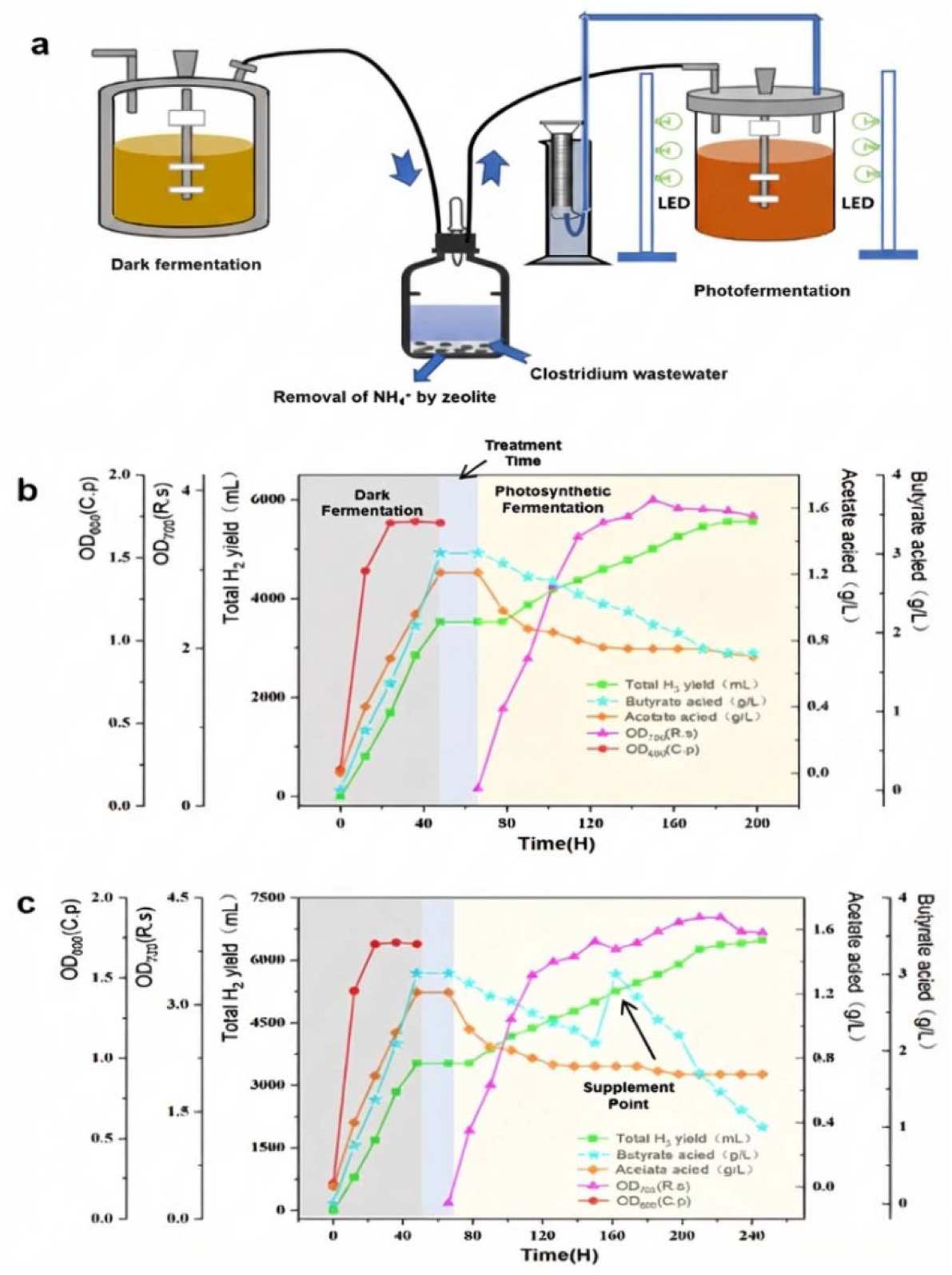
Changes in relevant parameters over time during the light-dark two-stage coupled H_2_ production process in a 5L bioreactor. (a) Schematic diagram of two-stage hydrogen production and fermentation with light-dark coupling. (b) Changes in relevant parameters over time during the two-stage coupled hydrogen production process without introducing the feeding process, where dark fermentation lasts for 48 h, followed by a treatment period of 18 h. The treated wastewater is used for hydrogen production by photosynthetic bacteria with a cycle of 132 h. (c) Changes in relevant parameters over time during the two-stage coupled hydrogen production process after introducing the feeding fermentation process, where dark fermentation lasts for 48 h, the treatment period is 18 h, and photo-fermentation by photosynthetic bacteria lasts for 168 h. The feed liquid is continuously supplied at 84 h of photo-fermentation (corresponding to 172 h of the total H_2_ production process), the butyric acid concentration is controlled at 3.0 g/L

Upon analyzing the data, the photo-fermentation stage demonstrated a significant reduction in dark fermentation products. Acetic acid levels decreased from 1.2 g/L to 0.61 g/L (representing a 50.1% removal rate), and butyric acid levels were cut from 3 g/L to 1.76 g/L (a 42.33% reduction). In comparison to a standalone dark fermentation system, which produced a cumulative 3526.6 mL of H2 over 72 hours, the integrated process boosted H_2_ production by an additional 2030.5 mL through photo-fermentation, culminating in a total H_2_ production of 5557.1 mL—an increase of 29.7%. This outcome underscores the synergistic potential of secondary organic acid conversion. Kinetic analysis revealed that dark fermentation’s peak H2 production occurred within the first 48 hours (comprising 97.4% of the total), whereas photo-fermentation exhibited higher H2 production between 96-160 hours. This two-stage process achieved metabolic complementarity by aligning production timelines. The remaining butyric acid level at the conclusion of photo-fermentation was 1.74 g/L, which was below the feeding threshold of 2 g/L. At this juncture, the photo-fermentation process was near its metabolic limit, suggesting that a combined feeding strategy could enhance the overall H_2_ production of the system. Adhering to the feeding strategy outlined in section 4.2, the feed liquid was introduced continuously starting at the 84th hour of photo-fermentation (corresponding to the 172nd hour of the total H_2_ production process), maintaining a butyric acid concentration of 3.0 g/L. Subsequently, the residual substrate concentration was monitored, and additional feeding was performed when the butyric acid concentration fell below 2.0 g/L. The final fermentation results are detailed in **Fig 5c**.

Without additional substrate, the control group finally had a total H_2_ production of 5557.1 mL, while the feeding group, through precise concentration monitoring and feeding control, continuously increased the H_2_ production to 6480.36 mL at 240 h, with a total output increase of 16.4%, effectively extending the H_2_ production cycle to 2.3 times that of the non-feeding group. Compared with the single dark fermentation process (total H_2_ production of 3372.4 mL), the H_2_ production of the coupled process increased by 80%. Compared with the single photo-fermentation feeding process (total H_2_ production of 3260.36 mL), the H_2_ production of the coupled process increased by 87%. Table 1 summarizes recent advances in dark-light coupled hydrogen production processes. Among systems with reactor volumes exceeding 1 L, the maximum H□ yield and average H□ production rate (L/d) achieved in this study are more representative, offering a new paradigm and perspective for the industrialization of H_2_ production.

**Table 1.** Advances in dark-light coupled biohydrogen production processes.

| Strains | Production Scale | Maximum H<sub>2</sub> Yield | Average H<sub>2</sub> production rate (L/d) | Ref |
| --- | --- | --- | --- | --- |
| *Clostridium butyricum* and *Rhodospseudomonas faecalis* RLD-53 | 100 mL | 5.374 mol-H<sub>2</sub>/mol-glucose | 0.0589 | (Bing-Feng et al., 2010) |
| *R. sphaeroides* NMBL-01, *Bacillus firmus* NMBL-03 | 120 mL batch reactor | 1510 mL/L | 0.0274 | (Anjana et al., 2020) |
| *Marichromatium purpuratum* LC83, *Pantoea agglomerans* BH-18 | 250 mL serum bottle | 1240 mL/L | 0.0150 | (Zhu et al., 2016) |
| *Enterobacter cloacae* DM 11, *R. sphaeroides* O.U. 001 | 300 mL batch reactors | 1238 mL/L | 0.0125 | (Nath & Das, 2009) |
| Microflora and *R. sphaeroides* | 600 mL(DF) and 250 mL(PF) | 713.6 ± 44.1 mL H<sub>2</sub>/g-COD | 0.0278 | (Liu et al., 2015) |
| *Escherichia coli*, *Clostridium acetobutylicum*, *R. capsulatus* | 1 L fermenter | 5.5 mol H<sub>2</sub>/mol glucose | 0.0346 | (Morsy et al., 2019) |
| *C. acetobutylicum*, *R. sphaeroides* | 1 L reactor | 3.23 L H<sub>2</sub>/L<sub>med</sub> | 0.0352 | (Zagrodnik & Waniecki, 2016) |
| *C. butyricum* LS2, *R. palustris* | 2 L CSTR | 3.064 ml H<sub>2</sub>/mL POME | 0.0206 | (Mishra et al., 2016) |
| *C. pasteurianum*(DF), *R. sphaeroides* ZX-5(PF) | 10 L bioreactor | 5.654 mol H<sub>2</sub>/mol glucose ( 275 3.6 mL/L ) | 0.0648 | This study |

However, according to the theoretical reaction for anaerobic fermentation of glucose to produce H_2_ (C_6_H_12_O_6_ + 6H_2_O → 12H_2_ + 6CO_2_), the theoretical H_2_ production from 30 grams of glucose (with a molecular weight of 180 g/mol, equivalent to 0.1667 moles) is 2.0004 moles of H_2_. This corresponds to a volume of 44810 milliliters under standard temperature and pressure (STP); however, the actual hydrogen production in the experiment was 6480.36 milliliters. Therefore, the conversion rate of glucose to hydrogen in a single batch was 14.46%. This conversion rate is lower than the optimized levels reported in the literature (for example, 4.0 moles of H_2_ per mole of glucose, equivalent to an 80.21% conversion rate, indicating that there is significant potential for improvement in the current process. Potential areas for improvement include optimizing fermentation conditions, such as pH, temperature, and microbial strain adaptation, to enhance the overall H_2_ yield. Furthermore, exploring the precise regulation of substrate feeding and optimizing the light/dark cycles during the photo-fermentation phase could help maximize H_2_ production. The current results highlight the promising potential of the coupled process but also emphasize the need for further optimization and scale-up efforts to achieve higher conversion efficiencies. Future research should concentrate on improving the metabolic pathways of the strains used, as well as refining bioreactor design and operation strategies to enhance the economic feasibility of large-scale biohydrogen production.

## 4. Conclusion

This study underscores the potential of integrating dark and photo-fermentation processes for the production of biohydrogen using *R. sphaeroides* ZX-5 and *C. pasteurianum* DSM 525. The 5L coupled system produced 6480.36 mL of H_2_, marking an 80% increase over single-stage dark fermentation, along with significant degradation of acetic acid (50.10%) and butyric acid (42.33%). The optimization of the photo-fermentation bioreactor process was crucial in enhancing H_2_ yield and optimizing organic waste utilization. In summary, the dark-light coupled process demonstrates considerable promise for sustainable H_2_ production and clean energy generation.

E-supplementary data of this word can be found in online version of the paper.

## Appendix A. Supplementary data

Fig. S1 to S6 Table S1 to S3

## Credit Author Statement

L.Z., G.-Y.T., H.F. and X.Z. conceived and designed the project. W.W., X.Z., Y.C. and L.Y. performed the experiments. W.W., X.Z. and Y.C. analyzed the results. W.W. and X.Z. wrote the manuscript with contributions from all co-authors.

## Acknowledgments

The National Research and Development Program Project (2020YFA0907304) and the National Natural Science Foundation of China (32370064 and 31870040) are acknowledged.

