## Supplementary Materials for "Enhanced Hydrogen Production Through Two-Stage Fermentation Coupling *Clostridium pasteurianum* and *Rhodobacter sphaeroides*"

|  | **Designation** | **Concentration(g·L^−1^)** |
| --- | --- | --- |
| **Substrate** | **DL-malate** | **4** |
| **Trace Element Solutions**  **(1 mL/L)** | **MnSO_4_·4H_2_O (g/L)** | **2.1** |
|  | **H_3_BO_3_ (g/L)** | **2.8** |
|  | **Cu(NO_3_)_2_·7H_2_O (g/L)** | **0.04** |
|  | **ZnSO_4_·7H_2_O (g/L)** | **0.24** |
|  | **Na_2_MoO_4_·2H_2_O** | **0.75** |
| **Metal elements** | **CaCl_2_·2H_2_O** | **0.075** |
|  | **MgSO_4_·7H_2_O** | **0.2** |
|  | **FeSO_4_·7H_2_O** | **0.0118** |
|  | **Ethylenediaminetetraacetic Acid Disodium Salt** | **0.04** |
| **Phosphate Buffered Solution (PBS, 0.2 M, PH=7.0)(7.5 mL/L)** | | |
| **Vitamin solution** | **Nicotinic Acid (g/L)** | **10** |
|  | **Thiamine (Vitamin B_1_)（g/L）** | **5** |
|  | **D-biotin（g/L）** | **0.1** |

**Table S1  Composition of RCVBN Medium**

**Table S2  Composition of MSG liquid medium**

|  | **Designation** | **Concentration(g·L^−1^)** |
| --- | --- | --- |
| **Trace Element Formulation** | **FeCl_2_·4H_2_O** | **2.00** |
|  | **ZnCl_2_** | **0.05** |
|  | **MnCl_2_·4H_2_O** | **0.05** |
|  | **CuCl_2_·2H_2_O** | **0.03** |
|  | **(NH_4_)_6_Mn7O_2_·4H_2_O** | **0.05** |
|  | **AlCl_3_** | **0.05** |
|  | **CoCl_3_·6H_2_O** | **0.20** |
|  | **Saturated H_3_BO_3_ Solution** | **1.0 mL** |
|  | **HCl** | **1.0 mL** |
| **Vitamin Solution Formula** | **Biotin** | **0.002** |
|  | **Folic acid** | **0.002** |
|  | **Pyridoxine-HCl** | **0.010** |
|  | **Riboflavin** | **0.005** |
|  | **Thiamine-HCl** | **0.005** |
|  | **Niacin** | **0.005** |
|  | **Cyanocobalamin** | **0.005** |
|  | **P-aminobenzoic acid** | **0.005** |
|  | **Pantothenic acid** | **0.005** |
| **Mineral Salt Solution Formula** | **NH_4_Cl** | **6.0** |
|  | **NaCl** | **6.0** |
|  | **CaCl_2_·2H_2_O** | **0.2** |
|  | **MgCl_2_·6H_2_O** | **2.0** |

**Table S3  Table S2  Composition of MSG liquid medium**

|  | **Designation** | **Concentration** |
| --- | --- | --- |
| **Substrate** | **Glucose(g/L)** | **1.274** |
| **VFAs** | **Butyric acid (g/L)** | **3.123** |
|  | **Acetic acid (g/L)** | **1.28** |
|  | **Succinic acid (g/L)** | **0.007** |
|  | **Formic acid (g/L)** | **0.003** |
|  | **Oxalic acid (g/L)** | **0** |
|  | **Malic acid (g/L)** | **0** |
| **Metal elements** | **Ca (mg/L)** | **10** |
|  | **Mg (mg/L)** | **5.6** |
|  | **Na (mg/L)** | **2700** |
|  | **Zn (mg/L)** | **0.93** |
| **N source** | **Ammonia nitrogen (mg/L)** | **6** |
|  | **Total nitrogen (mg/L)** | **144** |
| **COD** | **COD (mg/L)** | **10190** |

**(Note:** **VFAs stand for Volatile Fatty Acids, COD stand for Chemical Oxygen Demand)**

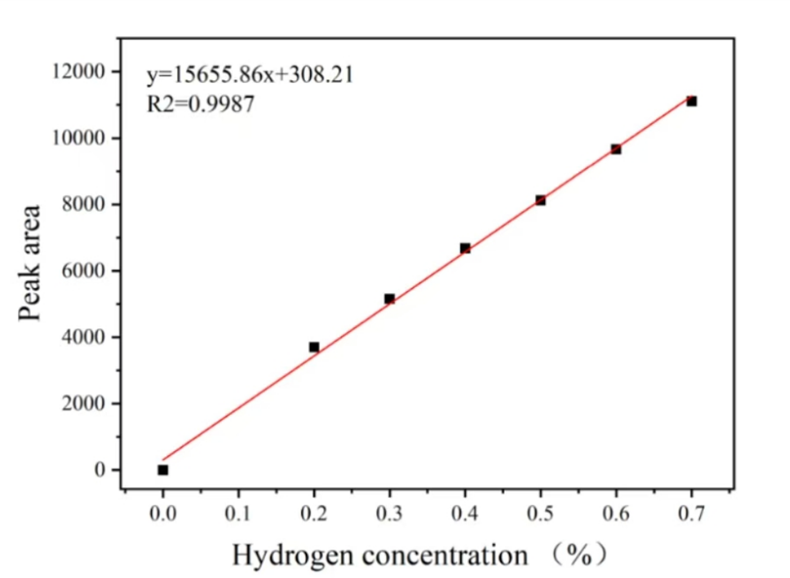

**Fig. S1 Hydrogen determination standard curve. The X-axis represents peak area, and the Y-axis represents the corresponding hydrogen concentration (g/L)**

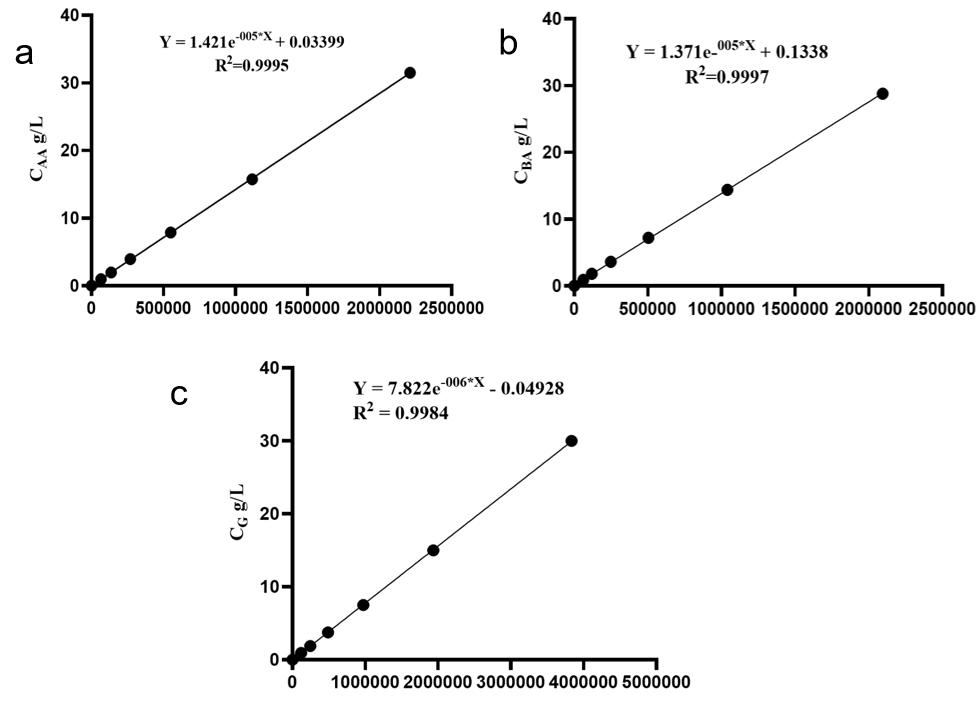

**Fig. S2 Standard curves for carbon source quantification. (a) Standard curve for acetate quantification. (b) Standard curve for butyrate quantification. (c) Standard curve for glucose quantification. The X-axis represents peak area, and the Y-axis represents the corresponding substrate concentration (g/L).**

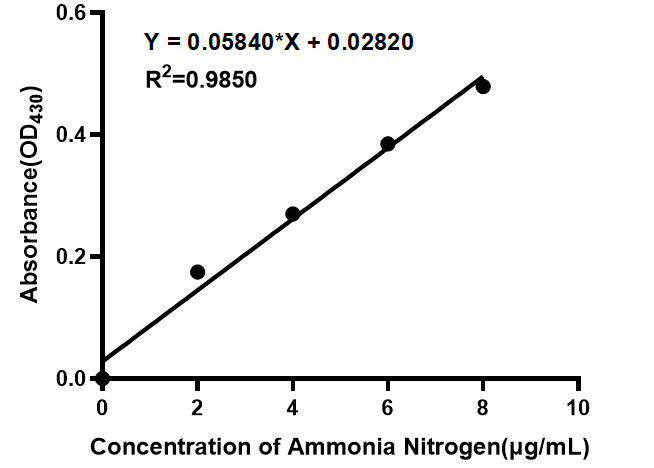

**Fig. S3 Standard curve for ammonia nitrogen determination. The X-axis represents concentration of ammonia nitrogen(μg/mL), and the Y-axis represents the absorbance.**

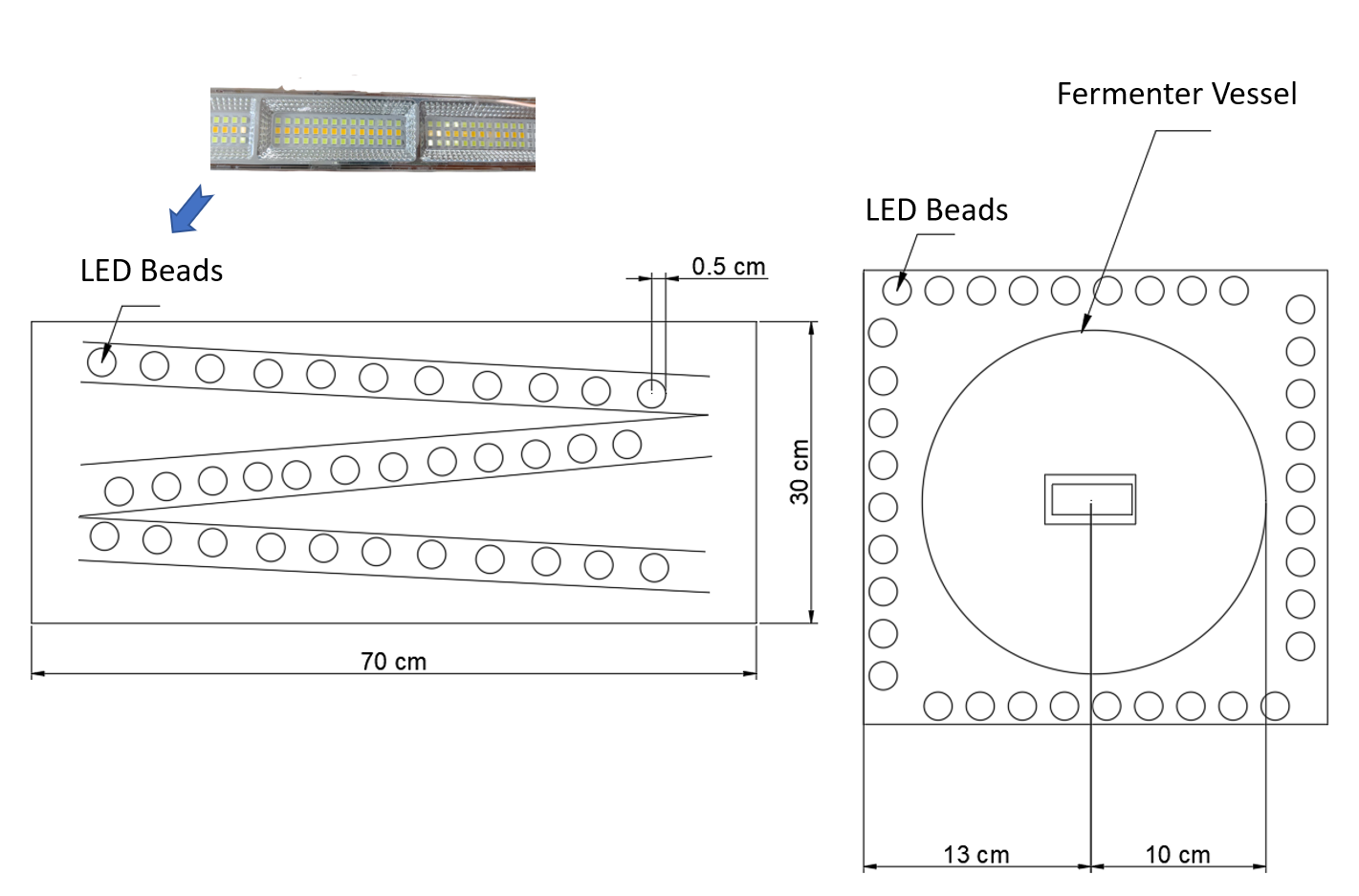

**Fig. S4 Integrated Photo-bioreactor with Orbital Light Source and Conventional Fermenter. The left panel details a rectangular LED array (70 cm × 30 cm) featuring four parallel rows of tightly spaced LED beads, each row separated by 0.5 cm. This configuration ensures uniform light distribution for photobioreactor applications. The right panel shows a top-down view of the fermenter vessel (10 cm width) with a central circular region (13 cm diameter). LED beads are arranged concentrically around this core. A small rectangular port at the base suggests sampling or monitoring access.**

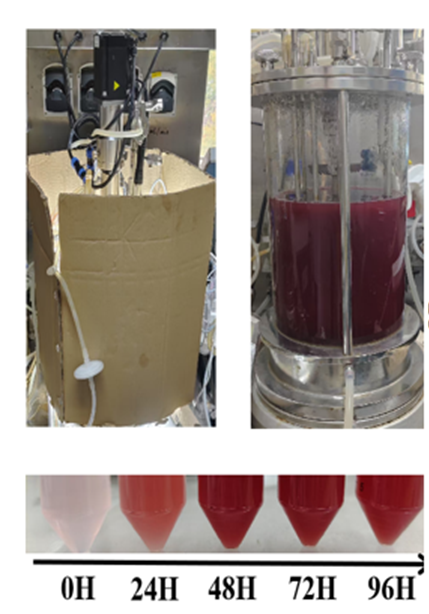

**Fig. S5 Hydrogen Production by Photo-fermentation in a 5L Reactor. The bottom section displays a time-series of conical flasks (0H to 96H), showing a progressive deepening of red hue, indicating microbial metabolic activity and hydrogen yield over 96 hours. The setup includes integrated pipelines and sensors for process monitoring.**

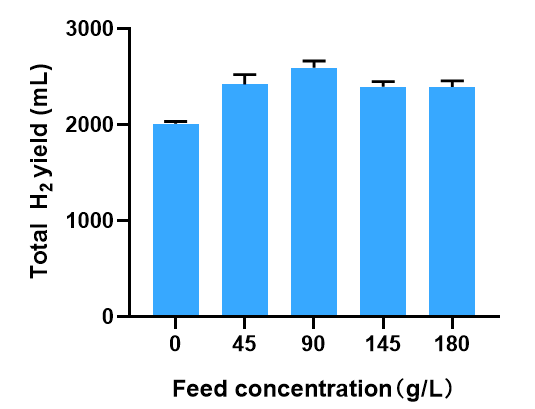

**Fig. S6  Effect of feed concentration on the growth of ZX-5 and hydrogen production.**
